# Cardiomyocyte-specific Srsf3 deletion reveals a mitochondrial regulatory role

**DOI:** 10.1101/2020.07.03.186999

**Authors:** Audrey-Ann Dumont, Lauralyne Dumont, Delong Zhou, Hugo Giguère, Chantal Pileggi, Mary-Ellen Harper, Denis P Blondin, Michelle S Scott, Mannix Auger-Messier

## Abstract

Srsf3 was recently reported as being necessary to preserve RNA stability via an mTOR mechanism in a cardiac mouse model in adulthood. Here, we demonstrate the link between Srsf3 and mitochondrial integrity in an embryonic cardiomyocyte-specific Srsf3 conditional knockout (cKO) mouse model. Fifteen-day-old Srsf3 cKO mice showed dramatically reduced (below 50%) survival and reduced left ventricular systolic performance, and histological analysis of these hearts revealed a significant increase in cardiomyocyte size, confirming the severe remodelling induced by Srsf3 deletion. RNA-seq analysis of the hearts of 5-day-old Srsf3 cKO mice revealed early changes in expression levels and alternative splicing of several transcripts related to mitochondrial integrity and oxidative phosphorylation. Likewise, the levels of several protein complexes of the electron transport chain decreased, and mitochondrial complex I-driven respiration of permeabilized cardiac muscle fibres from the left ventricle was impaired. Furthermore, transmission electron microscopy analysis showed disordered mitochondrial length and cristae structure. Together with its indispensable role in the physiological maintenance of mouse hearts, these results highlight the previously unrecognized function of Srsf3 in regulating mitochondrial integrity.

## Introduction

In the past three decades, splicing factors have been widely investigated, and some of them have been shown to belong to the serine/arginine-rich splicing factor (Srsf) family, members of which have a well-conserved RS domain throughout higher eukaryotic species (Boucher *et al*, 2001). This family is composed of 12 canonical members and several functions relating to a variety of biological processes beyond RNA splicing are attributed to these proteins (Howard & Sanford, 2015; Jeong, 2017). Specifically, Srsf3 is associated with the cell cycle (Kurokawa *et al*, 2014), cell proliferation (Ajiro *et al*, 2016; Jia *et al*, 2010; Song *et al*, 2019), apoptosis (Kim *et al*, 2014), glucose and lipid metabolism (Sen *et al*, 2013), and other key cellular processes (Howard & Sanford, 2015) in different organs such as the liver (Kumar *et al*, 2019; Sen *et al*., 2013), brain (Boutej *et al*, 2017; Song *et al*., 2019; Wong *et al*, 2012), and skeletal muscles (Rawcliffe *et al*, 2016). Furthermore, two recent articles reported splicing activity for Srsf3 in cardiac models. The first article described several unstable mRNAs involved in cardiac contraction due to a lack of 5’ capping in Srsf3 conditional knockout (cKO) mice through an Srsf3-mTOR-eIF4E mechanism (Ortiz-Sanchez *et al*, 2019). The second study revealed the capacity of p38 MAPK to modulate the splicing activity of Srsf3 in cultured primary neonatal rat ventricular myocytes (Dumont *et al*, 2019).

Abnormal RNA splicing is an important contributor to the development of cardiomyopathies (Beqqali, 2018; Kim *et al*, 2018; Kong *et al*, 2010; Lara-Pezzi *et al*, 2013; Lee *et al*, 2011). In fact, many studies have been performed to understand the impact of distinct splicing factors on mRNA expression and splicing from a cardiac perspective via the use of human heart samples and cardiac mouse models. In some cases, the development of cardiomyopathy affected the expression of splicing factors and their mRNA regulating functions. For example, RBM25/LUC7L3 and RBFox1 had their expression altered in human heart failure, which, in turn, affected the splicing of important cardiac mRNAs such as SCN5A and different Mef2 isoforms (Gao *et al*, 2016; Gao *et al*, 2011). Furthermore, several cardiomyocyte-specific cKO of splicing factors in mice such as hnRNPU (Ye *et al*, 2015), Srsf1 (Xu *et al*, 2005), Srsf2 (Ding *et al*, 2004), RBFox2 (Wei *et al*, 2015), and RBM24 (Liu *et al*, 2019) led to the development of dilated cardiomyopathy at different ages. These depletions impair the splicing and expression of critical mRNAs for normal heart function. Most splicing factor studies, including three studies examining Srsf3 (Ortiz-Sanchez *et al*., 2019), Srsf10 (Feng *et al*, 2009) and RBM20 (Guo *et al*, 2012), reported several mRNA alterations related to sarcomeric and ion handling/channel genes, which are essential for adequate heart contraction and relaxation processes (de Groot *et al*, 2020; Ding *et al*., 2004; Gao *et al*., 2011; Xu *et al*., 2005; Yang *et al*, 2014).

Another primordial process needed to sustain the high energy demand of cardiac contraction-relaxation cycles is adenosine triphosphate (ATP) production by the mitochondrial oxidative phosphorylation system. The heart is the highest consumer of ATP as a percentage of human body weight (Neubauer, 2007). It is also the organ containing the highest mitochondrial content per unit of cellular volume (Brown *et al*, 2017). There are a few examples of splicing events on specific mRNAs related to the different processes regulating ATP production in different organs. For example, PTB (hnRNPI) together with hnRNPA1 and hnRNPA2 modulates the isoform change of PKM1 to PKM2 (pyruvate kinase M) through the splicing event of two mutually exclusive exons, which partially modulates the switch from oxidative phosphorylation to aerobic glycolysis in cancer cells (Chen *et al*, 2010b; David *et al*, 2010). Aberrant splicing of transcripts of the iron-sulfur cluster assembly gene (*ISCU*) is produced by a mutation in a cryptic intronic splicing site, leading to a nonfunctional protein, and is partly modulated by SRSF3 in human myoblasts (Rawcliffe *et al*., 2016). The ISCU has a role as a scaffold protein, and the mutation ultimately affects the respiratory chain complexes and the Krebs cycle in the skeletal muscles of individuals affected by hereditary myopathy with lactic acidosis.

Studies focusing on cardiac models also reported a change in mRNA linked to the generation of ATP and mitochondrial function. Indeed, SF3B1 mediates the A-to-C isoform change of ketohexokinase, leading to ATP depletion in mouse models of pathological cardiac hypertrophy (Mirtschink *et al*, 2015). Mitofusin 2, involved in mitochondrial fusion (Dorn, 2020), and Slc25a3, the mitochondrial inorganic phosphate carrier (Mayr *et al*, 2007), are alternatively spliced by RBM24 (Liu *et al*., 2019). As these are examples of only a few specific cardiac genes related to ATP production, they are very important and suggest that splicing factors may have a meaningful impact on energy production in the heart.

Here, we report the impact of a new cardiomyocyte-specific cKO of the splicing factor Srsf3 in neonatal mice. Conditional depletion of Srsf3 in the mouse cardiomyocytes led to a significantly increased cell size accompanied by systolic dysfunction. RNA-seq performed on the hearts of these mice revealed an unexpected cluster of mRNAs related to oxidative phosphorylation. With a variety of analyses, cardiac mitochondria were shown to have several irregularities, suggesting impairment of their function to produce energy in the absence of Srsf3.

## Results and Discussion

### Srsf3 deletion impairs cardiac function and leads to rapid death

To study the role of Srsf3 in the heart, we generated a cardiomyocyte-specific Srsf3 cKO mouse model by breeding Srsf3-floxed (Sen *et al*., 2013) and βMHC-Cre (Parsons *et al*, 2004) transgenic mice, the latter of which expresses Cre recombinase in cardiomyocytes driven by the 5.6 kb mouse βMHC promoter during embryogenesis. At birth, the Mendelian ratio was relatively normal, with approximately 22% for Srsf3^Fl/+-βMHC-cre^ and 21% for Srsf3^Fl/Fl-βMHC-Cre^ living progeny, given the expected 25% probability in theory for each of these genotypes (Fig 1A). Such an insubstantial reduction in birth rate is in sharp contrast with a similar cardiomyocyte-specific Srsf3 cKO recently reported (Ortiz-Sanchez *et al*., 2019). Indeed, this previous study failed to generate any homozygous Srsf3-floxed offspring expressing Cre recombinase under distinct Nkx2.5 and αMHC cardiac-specific promoters at birth. The onset of expression of the Cre recombinase under the regulation of these promoters could explain why such marked survival difference is observed between the different cardiomyocyte-specific Srsf3-floxed crosses. Lineage tracing approaches revealed that Cre-mediated embryonic recombination is detectable at as early as E8.0-8.5 in either Nkx2.5-Cre knock-in (Zhou *et al*, 2008) or αMHC-Cre (McFadden *et al*, 2005) transgenic mice. However, efficient recombination of the floxed gene in cardiomyocytes from βMHC-Cre crosses is barely established at E12.5 and starts only at later developmental stages (*i*.*e*., E17.5) of the heart (Oka *et al*, 2006). Nonetheless, the fact that Srsf3^Fl/Fl-βMHC-Cre^ mice survive birth, unlike Srsf3^Nkx2.5-Cre^ and Srsf3^αMHC-Cre^ mice, provided us with the opportunity to investigate the role of this RNA-binding protein in the heart of neonates.

**Figure 1.**
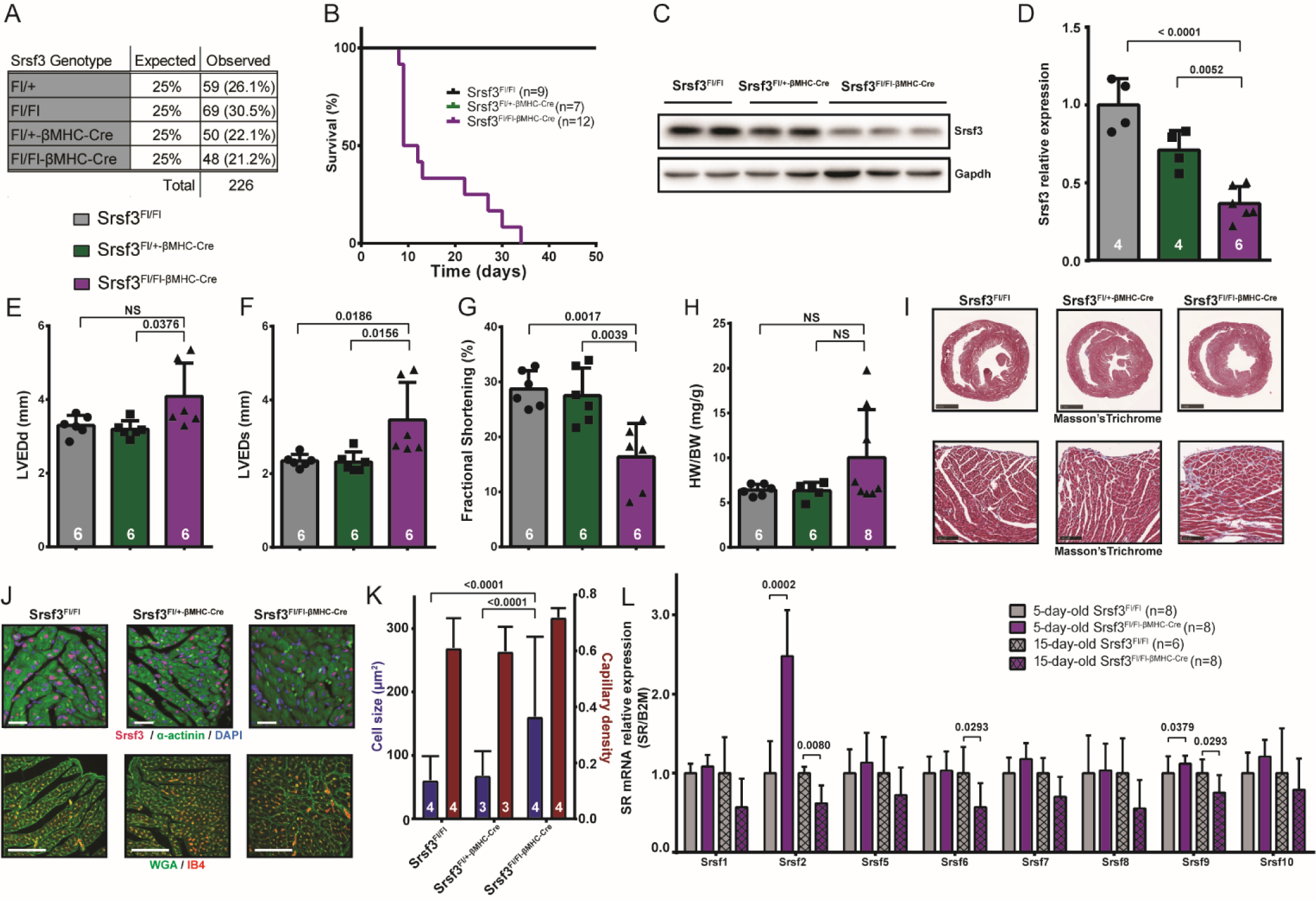
Cardiac deletion of Srsf3 during embryogenesis impairs heart function. **A)** Table of the expected and observed Mendelian ratio of Srsf3-floxed +/- βMHC-Cre offspring from the mating of Srsf3^Fl/+-βMHC-cre^ and Srsf3^Fl/Fl^ mice. **B)** Representation of the mouse survival curve for each indicated genotype. Note that lines corresponding to Srsf3^Fl/Fl^ and Srsf3^Fl/+-βMHC-cre^ mice are superimposed. **C, D)** Representative western blots revealing the expression of the Srsf3 protein and Gapdh loading control from whole heart lysates of the indicated genotypes. Srsf3 relative expression normalized to Gapdh and quantified from the western blots. **E-G)** Measurements of the left ventricular end-diastolic diameter (LVEDd), left ventricular end-systolic diameter (LVEDs), and heart contractility represented by the fractional shortening obtained by echography from 15-day-old mice. **H)** Gravimetric analysis of the heart weight-to-body weight (HW/BW) ratio from 15-day-old mice of each indicated genotype. **I)** Heart transverse slices stained with Masson’s trichrome (scale bars: 100 μm, upper panels; 1 mm, lower panels) from 15-day-old mice. **J, K)** Histological images of transverse sections of the heart from each indicated genotype and stained with Srsf3 (red) reveal its colocalization at the DAPI-labelled nucleus (blue) relative to the sarcomere’s immunostained α-actinin (green) (scale bar: 10 μm, upper panels). Immunostained wheat germ agglutinin (WGA, green) and isolectin B4 (IB4, red) to identify the plasma membrane and endothelial cells from transverse sections of the heart from each indicated genotype (scale bar: 100 μm, lower panels). The pictures of WGA/IB4-stained sections were used to measure the cell surface area and the density of capillaries per cardiomyocyte, which are shown as blue and red bars, respectively. **L)** RT-qPCR of the different members of the SR protein family from 5- and 15-day-old Srsf3^Fl/Fl^ and Srsf3^Fl/Fl-βMHC-cre^ mice.

Notably, half of the Srsf3^Fl/Fl-βMHC-Cre^ mice died within ten days after birth, and none survived beyond five weeks, while both Srsf3^Fl/Fl^ and Srsf3^Fl/+-βMHC-cre^ control mice aged normally (Fig 1B). This observation is in contrast with the longer life expectancy of Srsf1 and Srsf2 cKO mouse models following their invalidation during embryonic development with the ventricular myocyte restricted Mlc2v-Cre transgene (Chen *et al*, 1998; Ding *et al*., 2004; Xu *et al*., 2005). Indeed, none of the Srsf1 cKO mice died before 5 weeks of age, whereas more than 80% of the Srsf2 cKO mice were still alive 20 weeks after birth. We analysed the relative Srsf3 expression in whole ventricle lysates from 15-day-old mice, and compared to the results for the control Srsf3^Fl/Fl^ mice, we observed progressively diminished expression, averaging 71.1% and 36.8% for Srsf3^Fl/+-βMHC-cre^ and Srsf3^Fl/Fl-βMHC-Cre^ mice, respectively (Fig 1C-D). Echocardiographic analysis hinted at the eccentric remodelling of Srsf3^Fl/Fl-βMHC-Cre^ mice, as their left ventricular end diastolic diameter (LVEDd) tended to increase, although not significantly (Fig 1E). However, the contractile performance from these same mice was clearly diminished, as indicated by the 1.5-fold enlarged left ventricular end-systolic diameter (LVEDs) (Fig 1F), the marked (48%) reduction in fractional shortening (Fig 1G), and the significantly lower cardiac output (Appendix Table S1) compared to control Srsf3^Fl/Fl^ mice. The amplitude of systolic dysfunction developed in juvenile Srsf3 cKO mice is comparable to the overall cardiac effect of depleting other SR proteins, Srsf1 and Srsf2 (Ding *et al*., 2004; Xu *et al*., 2005), although this effect developed at very different ages, suggesting different underlying mechanisms.

Although the majority of 15-day-old Srsf3 cKO mice showed no significant increase in their heart weight- to-body weight (HW/BW) ratio, some animals clearly transitioned faster towards heart failure, which was accompanied by an important increase in the HW/BW ratio (Fig 1H), suggesting an incomplete penetrance or a variable expressivity of Srsf3 loss in cardiomyocytes. Accordingly, a heart cross-section stained with Masson’s trichrome revealed no obvious sign of gross morphological defects in most of the Srsf3 cKO mice analysed (Fig 1I, upper panels), although few showed clear dilated left ventricles, as observed in older Srsf1 and Srsf2 cKO mice (Ding *et al*., 2004; Xu *et al*., 2005). These two mouse models developed dilated cardiomyopathy within six and five weeks, respectively. Nevertheless, a perceptible onset of interstitial fibrosis was observed in the Srsf3^Fl/Fl-βMHC-Cre^ mice (Fig 1I, lower panels). The hepatocyte-specific Srsf3 cKO mouse model also revealed an increase in fibrosis in the liver upon deletion of this RNA-binding protein (Sen *et al*, 2015).

The residual expression of Srsf3 that we observed (Fig 1C-D) can likely be attributed to other cell types in the heart, specifically fibroblasts and endothelial cells. This is further supported by the immunostaining of Srsf3 performed on heart histologic sections that validated the cKO of nearly all cardiomyocytes in Srsf3^Fl/Fl-βMHC-Cre^ mice (Fig 1J, upper panels). Endothelial cells from blood vessels were stained with isolectin B4 (Fig 1J, lower panels) to quantify the capillary density between blood vessels and cardiomyocytes (Fig 1K, red bars). All three genotypes of mice demonstrated a similar proportion of capillaries per cardiomyocyte. The cell size of cardiomyocytes was assessed by membrane staining with wheat germ agglutinin (WGA) of a transverse heart section (Fig 1J, lower panels) followed by analysis of the cardiomyocyte surface area (Fig 1K, blue bars). The Srsf3^Fl/Fl-βMHC-Cre^ mice demonstrated significant cellular hypertrophy, whereas heart hypertrophy was not observed, suggesting a decreased number of cardiomyocytes in the whole heart. This is surprising given the capacity of cardiomyocytes to proliferate and compensate when cell loss occurs during foetal heart development (Cui *et al*, 2018; Sturzu *et al*, 2015). On the other hand, Srsf3 is well known to be linked to cell proliferation (Ajiro *et al*., 2016; Jia *et al*., 2010; Song *et al*., 2019) and studies about the role of Srsf3 in the proliferation of cardiomyocytes during heart embryogenesis warrant more attention in the future.

Quantitative reverse transcription PCR (RT-qPCR) analysis was performed on genes known to be involved in cardiac hypertrophy; *Gata4* (Liang *et al*, 2001; Whitcomb *et al*, 2020), *Srf* (serum response factor) (Nelson *et al*, 2005; Zhang *et al*, 2001), *Myh6* (αMHC), *Myh7* (βMHC) (Rohini *et al*, 2010), and *Nppb* (Natriuretic peptide B) (Sergeeva & Christoffels, 2013). Five days after birth, only the expression of *Nppb* exhibited a gain (2.5-fold) in the Srsf3^Fl/Fl-βMHC-Cre^ mice, while *Myh7* presented an expression reduction (Appendix Fig S1A-E). Ten days later, Srf, αMHC and βMHC all displayed a decline in their mRNA levels, suggestive of more advanced cardiac molecular and pathological remodelling in the Srsf3 cKO mice. The decrease in all analysed mRNA at fifteen days could be explained by the degradation of mRNA induced by the Srsf3-mTOR mechanism found in the adult mouse model (Ortiz-Sanchez *et al*., 2019).

The expression of other SR protein family members was investigated given the role of this protein family in regulating each other’s expression (Anko *et al*, 2012). We analysed the mRNA transcripts of several SR proteins by RT-qPCR at two time points after birth: five and fifteen days (Fig 1L). A significant gain in *Srsf2* expression was detected at five days of age under our conditions, whereas the other SR protein transcripts analysed showed no change. In another study, a clip analysis with Srsf3 showed that it binds the mRNA of Srsf2 (Anko *et al*., 2012). Moreover, they observed a change in Srsf2 splicing under Srsf3 overexpression conditions; specifically, they observed an increase in an exon containing a poison cassette, leading to degradation through nonsense-mediated decay. This could explain the initial increase in Srsf2 mRNA in our condition given that Srsf3 is not there to promote the inclusion of the poison cassette. Interestingly, all the mRNA of the SR proteins studied at 15 days of age exhibited a reduction in expression, again possibly suggesting secondary effects of the Srsf3 deletion and a more advanced pathology.

With the characterization of the cardiac embryonic Srsf3 cKO mouse model, it would be interesting to compare it to an adult cKO model. A recent article reported the impaired heart function and abrupt death in adulthood of the cardiac tamoxifen-inducible Srsf3 cKO mouse model (Ortiz-Sanchez *et al*., 2019). Likewise, we obtained similar results in the characterization of our identical adult Srsf3^Fl/Fl-αMHC-MerCreMer^ cKO model. Seven days after a single tamoxifen injection, Srsf3 expression decreased by 81% compared to that in control mice (Appendix Fig S2A). For the mice with Srsf3 deprived cardiomyocytes, the heart contractility measured by echography revealed a fractional shortening of 24%, while the mean for control mice was approximately 32% (Appendix Fig S2B). Seven days after the sole tamoxifen injection, no hypertrophy or fibrosis was observed (Appendix Fig S2C). Nevertheless, a fast drop in survival was noticed between seven and eight days after the single tamoxifen injection (Appendix Fig S2D), consistent with a previous report (Ortiz-Sanchez *et al*., 2019). The deletion at the embryonic stage with βMHC-Cre seems more permissive than with tamoxifen-inducible αMHC-MerCreMer considering the life expectancy of each model after Srsf3 depletion.

### Transcript analysis reveals oxidative phosphorylation as a cluster affected by the cardiomyocyte-specific Srsf3 conditional knockout

A high-throughput RNA-seq analysis was performed to explore Srsf3 function in 5-day-old mouse hearts. The choice of this time-point was based on two aspects. First, we wanted a time-point after birth at which Srsf3 has as high an expression level as possible in the heart, and Srsf3 is known to be well expressed in this organ during the first week of life, after which its expression level decreases (Ortiz-Sanchez *et al*., 2019). Second, another of our concerns was to avoid a state of pathology that was too advanced to investigate the primary effect of the Srsf3 deletion. After the analysis of the RNA-seq results (Appendix Table S2), we obtained 1130 genes with a significant expression change (Fig 2A). The number of genes with altered expression was lower than that observed in other RNA-seq analyses by splicing factor knockout in cardiomyocytes, but this could be explained in part by the difference in sequencing depth reached between the studies (Liu *et al*., 2019; Ortiz-Sanchez *et al*., 2019). Also note that the measured level of cardiac Srsf3 expression in Srsf3^Fl/Fl-βMHC-Cre^ was 57% of that in Srsf3^Fl/Fl^ mice with this RNA-seq analysis (Appendix Fig S3).

**Figure 2.**
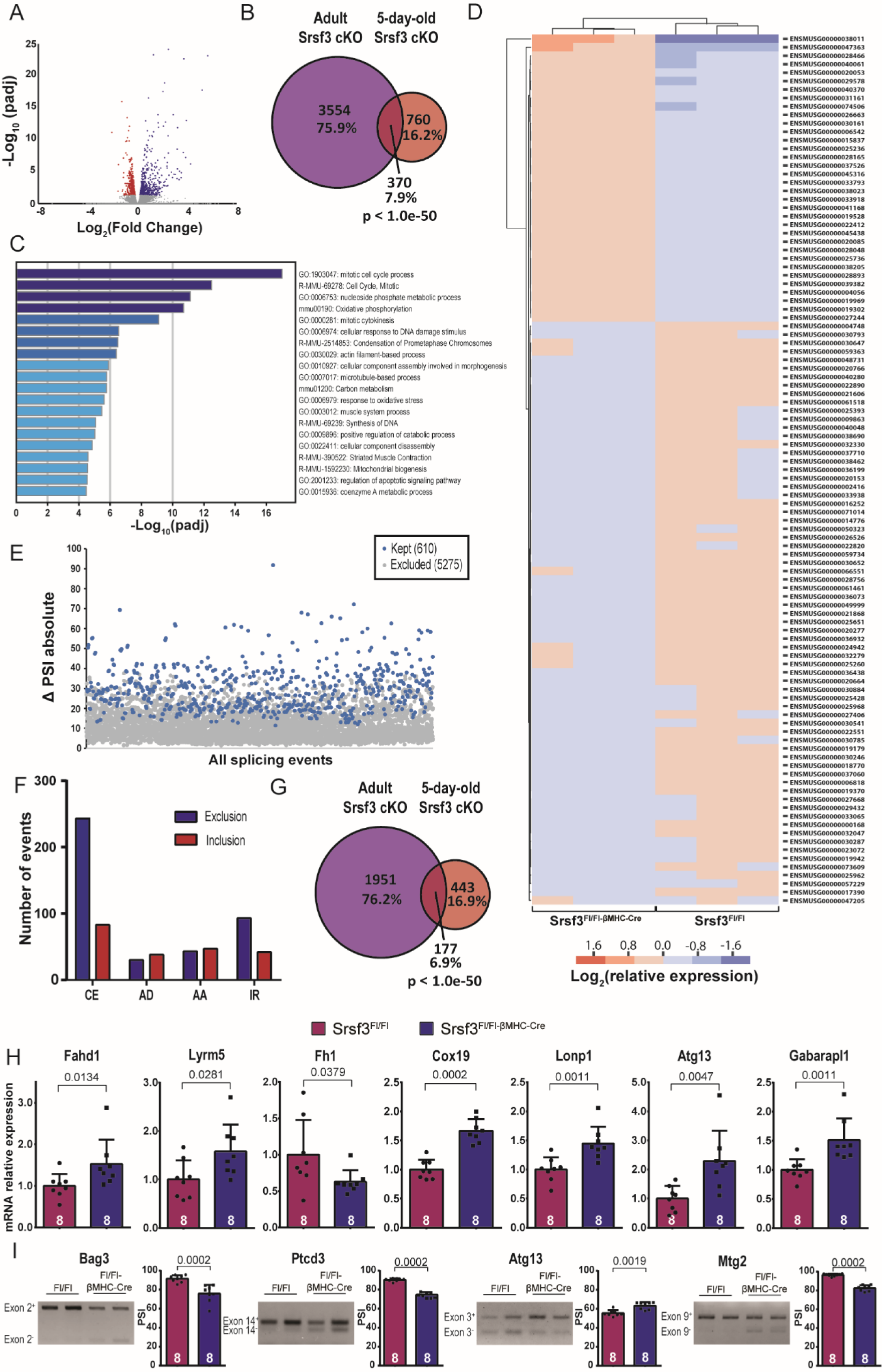
RNA-seq reveals the oxidative phosphorylation cellular process as a target of Srsf3 in the heart. **A)** Volcano plot of the mRNA expression level (blue and red dots are significant, whereas grey dots are not) from Srsf3^Fl/Fl-βMHC-Cre^ mice compared to Srsf3 Fl/Fl mice (5-day-old mice; n=3 for each group). **B)** Venn diagram comparison of the heart transcriptomes from the cardiomyocyte-specific Srsf3 cKO at early age (5-day-old Srsf3^Fl/Fl-βMHC-Cre^ mice) and adulthood (5 days after the initial tamoxifen injection of approximately 3-month-old Srsf3^Fl/Fl-αMHC-MerCreMer^ mice, as recently reported (Ortiz-Sanchez *et al*., 2019)). **C, D)** Metascape analysis performed on all mRNAs with a significant fold change in expression in Srsf3^Fl/Fl-βMHC-Cre^ mice. Each individual mRNA from the oxidative phosphorylation cluster identified is represented as a heat map. **E)** Scatterplot of all splicing events detected. Only the blue dots were retained (significant change in PSI higher than 10% compared to control mice). **F)** Number of each splicing event type (*i*.*e*., for the Srsf3 cKO mice compared to control mice). **G)** Venn diagram of all significant splicing events in a 5-day-old mouse heart compared to an adult mouse heart sample from E. Lara-Pezzi’s study (Ortiz-Sanchez *et al*., 2019). **H)** mRNA expression analysis by RT-qPCR of different mRNAs modulated by the absence of Srsf3 in cardiomyocytes implicated in oxidative phosphorylation. **I)** End-point PCR analysis of different spliced transcripts observed in the heart sample at an early age with the Srsf3 deletion model compared to the control.

Cardiomyocytes undergo a variety of changes between neonate and adult stages, one of which is modification of the transcription profile (Chen *et al*, 2004; Wang *et al*, 2018). With the available RNA-seq analysis performed on the heart from adult Srsf3 cKO mouse model (Ortiz-Sanchez *et al*., 2019), we were interested in determining how many genes with altered expression in our neonatal Srsf3 cKO model were shared with the adult model. A significant overlap (370 genes) between the two analyses was found, but several mRNAs remained unique to each study (Fig 2B). In neonate cardiomyocytes, important cellular processes are still occurring, such as cell division and proliferation, in contrast to the adult heart. Furthermore, Srsf3 was associated with the proliferation of cancer cells as a proto-oncogene (Ajiro *et al*., 2016; Jia *et al*., 2010), and also *in utero*, with the proliferative arrest of epicardial cells as shown from the study of proepicardium-specific Srsf3 cKO mice (Lupu, 2019). Enrichment analysis was performed via Metascape (Zhou *et al*, 2019) on the RNA-seq data, and interestingly, many gene ontology annotations implicated in proliferation, such as the mitotic cell cycle process and mitotic cytokinesis, were highlighted (Fig 2C). In cardiomyocytes, the latter cellular process demonstrated a defect during heart development in a GAS2L3 mouse deletion (Stopp *et al*, 2017) and we obtained similar conclusions in terms of the diminution of cardiomyocyte numbers in the heart as well as cellular hypertrophy. Two more unexpected gene ontology terms from the Reactome database (Jassal *et al*, 2020) interested us, namely, oxidative phosphorylation and mitochondrial biogenesis, due to the limited information between Srsf3 and those crucial processes for heart function. Oxidative phosphorylation takes place at the inner membrane of mitochondria and normally produces the majority of a cell’s ATP. A hierarchical clustering heatmap was generated to obtain a more exhaustive view of the expression difference of 103 mRNAs related to oxidative phosphorylation between the two mouse groups (Fig 2D). Knowing that splicing is an important function of Srsf3, we performed an alternative splicing analysis with Vast-tools and obtained 610 significant splicing events with a minimum threshold of 10% delta percent spliced in index (PSI) (Fig 2E). The most frequently observed alternative splicing-type event was the exon cassette (Fig 2F), which is the most common alternative splicing type (Kim *et al*, 2008). Moreover, the profile of all splicing events is akin to that observed in the Srsf3 cKO study conducted at adulthood (Ortiz-Sanchez *et al*., 2019). Furthermore, a significant fraction of the splicing events was preserved between the 5-day-old and adult Srsf3 cKO mice, while many splicing events were exclusive to each age group (Fig 2G). The overlap could be due to the similar focus of both studies, namely, the knockout of Srsf3 in the same cell type. However, the variation in splicing events between the studies may stem from the major transformations that occur during the transition from neonate to adult heart, including alternative splicing (Wang *et al*, 2016).

The RNA-seq analysis provided new insights into the transcriptome modulated by Srsf3, at the level of both expression and alternative splicing, in the mouse neonate heart. Several changes in mRNA expression (Appendix Fig S4A-G) implicated in the oxidative phosphorylation cluster were validated with heart samples of 5-day-old mice by RT-qPCR (Fig 2H). Some of the selected mRNAs have been associated with the Krebs cycle or respiratory chain complexes. This is the case for Fahd1 (Pircher *et al*, 2015; Weiss *et al*, 2018), Lyrm5 (Floyd *et al*, 2016; Pagliarini *et al*, 2008), and Fh1 (Tyrakis *et al*, 2017). Interestingly, a reduction in Fh1 leads to mitochondrial respiratory chain defects, particularly in the complex I and II (Tyrakis *et al*., 2017). Other validated mRNAs, such as Lonp1 or Atg13, have a mitochondrial protease function (Bota & Davies, 2016; Hoshino *et al*, 2014) or are linked to autophagy of the mitochondrion (Kaizuka & Mizushima, 2016), respectively. Furthermore, a few splicing events identified by the RNA-seq analysis (Appendix Fig S5A-D) were confirmed by alternative splicing PCR and analysed on an agarose gel (Fig 2I). Several mRNAs were also chosen (*i*.*e*., Bag3, Ptcd3, Atg13, and Mtg2) due to their association with mitochondria. Interestingly, Ptcd3 and Mtg2 are ribosomal proteins involved in mitochondrial protein translation (Datta *et al*, 2005; Davies *et al*, 2009; Kotani *et al*, 2013) and recently, the alternative splicing of a protein with similar function, the mitochondrial elongation factor GFM1, produced an alteration in several proteins serving in the assembly and activity of respiratory chain complexes suggestive of an oxidative phosphorylation defect (Simon *et al*, 2017). Moreover, several individuals with Bag3 mutations have been reported, and a few of them developed cardiomyopathy (Arimura *et al*, 2011; Fang *et al*, 2017; Lee *et al*, 2012; Norton *et al*, 2011; Selcen *et al*, 2009; Semmler *et al*, 2014). Some of the mutations are located on the excluded exon in the absence of Srsf3. A study in cardiomyocytes also revealed a quality control function of the mitochondria for the protein Bag3 (Tahrir *et al*, 2017). It would be interesting to investigate whether mutations in Bag3 can affect its splicing and protein structure and function in relation to the development of cardiomyopathies.

### Oxidative phosphorylation defects and loss of mitochondrial integrity in Srsf3 deficient cardiomyocytes

Splicing factors and mitochondria are not frequently associated with each other. However, high-throughput analysis helps to highlight these unexpected associations. Similar to our results, an RBM25 study using RNA-seq analysis coupled with gene ontology revealed mitochondria as one of the principal cellular components identified in HEK cells (Carlson *et al*, 2017). Another group reported alternative splicing of specific genes in the heart related to the mitochondria (*Nek1, Immt* and *Atp5c1*) via RASL-seq with an RBFox2 knockout model or a transverse aortic constriction (TAC) model (Wei *et al*., 2015). Based on our RNA-seq results suggesting that Srsf3 has an association with mitochondria, we decided to investigate the impact of cardiomyocyte-specific Srsf3 cKO on mitochondrial components and functionality in the cardiac muscle of mice.

Individuals with heart failure have marked alterations in cardiac tissues mitochondrial biogenesis. They display a reduction in mitochondrial DNA (mtDNA) content as well as hindered mtDNA replication and an increase in DNA oxidative damage (Karamanlidis *et al*, 2010). Srsf3^Fl/Fl-βMHC-Cre^ mice presented a decline of approximately 42% in the mtDNA content compared to Srsf3^Fl/Fl^ mice (Fig 3A). This reduction may be explained by the fact that Srsf3 has recently been linked to DNA repair via the regulation of genes related to this function. More precisely, a fraction of the genes were directly involved in mitochondrial DNA repair (He & Zhang, 2015). Interestingly, we also observed a gene association with the cellular response to DNA damage stimulus and synthesis of DNA in the Metascape results.

**Figure 3.**
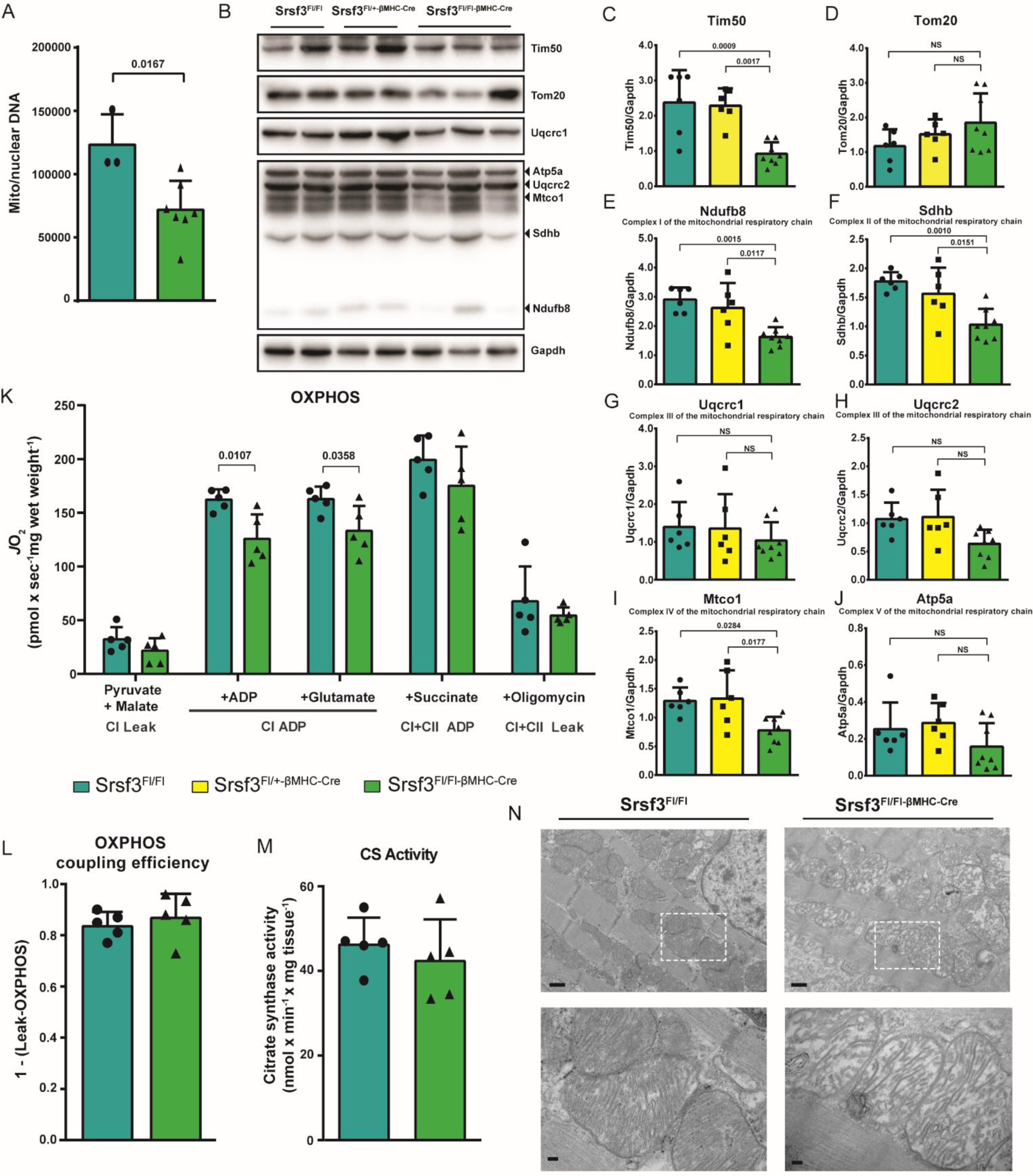
Srsf3 cKO in cardiomyocytes induces structural and functional mitochondrial changes. **A)** RT-qPCR analysis for relative quantification of mitochondrial DNA compared to nuclear DNA from Srsf3 cKO mouse heart tissues compared to samples from control mice. **B-J)** Representative western blots revealing the expression of discrete mitochondrial proteins and the Gapdh loading control from the whole heart lysates of the indicated genotypes. Bar graphs showing the quantified relative expression of each corresponding protein normalized to Gapdh. **K)** Mitochondrial respiration of permeabilized myocardial fibres from the left ventricular free wall of 8-day-old Srsf3^Fl/Fl^ and Srsf3^Fl/Fl-βMHC-Cre^ mice following the sequential addition of 2 mM malate (M) and 5 mM pyruvate (P) (adenylate-free leak respiration; complex I leak or CI_Leak_), followed by the addition of 4 mM ADP (D) and 10 mM glutamate (G) (complex I-supported respiration; CI_ADP_). Then, maximal oxidative phosphorylation (OXPHOS) capacity with convergent electron flow from complex I and complex II was reached with 10 mM succinate (S) (CI+CII_ADP_). ATP synthesis was then inhibited using 2.5 μM oligomycin to inhibit ATP synthase and induce non-phosphorylating leakage (CI+CII_Leak_). Finally, non-mitochondrial respiration was measured after complex III inhibition by 5 μM antimycin. **L)** OXPHOS coupling efficiency was calculated as 1 – (intrinsic CI_Leak_/ CI+CII_ADP_). **M)** Citrate synthase activity of homogenized left ventricular myocardial tissue. **N)** Representative transmission electron microscopy images focusing on the mitochondrion from the histological heart sections of 8-day-old Srsf3^Fl/Fl^ and Srsf3^Fl/Fl-βMHC-Cre^ mice at 7,000X (scale bar: 500 nm, upper panels) and 25,000X (scale bar: 100 nm, lower panels) magnifications.

The evaluation of the protein level localized to the mitochondria was another important aspect to the characterization of mitochondrial integrity. Translocases on the mitochondrial outer and inner membranes, Tom20 and Tim50, import proteins into mitochondria from the cytosol, and were analysed because they are modulated in heart pathologies. Tim50 is reduced in mice with hypertrophic hearts and in humans with dilated cardiomyopathy (Tang *et al*, 2017) while Tom 20 is diminished in ischaemic porcine hearts (Boengler *et al*, 2006). A reduction in Tim50 of approximately 60% was observed in the Srsf3 cKO mice compared to the control mice, whereas Tim20 showed no significant change (Fig 3B-3D). Given that the gene ontology analysis of the RNA-seq points towards oxidative phosphorylation, a western blot analysis was performed for some proteins associated with this function (Fig 3B, 3E-3J). Three of the six proteins investigated showed significant reductions of 40-44% in Srsf3 cKO mice, namely, Ndufb8 (Fig 3E), Sdhb (Fig 3F) and Mtco1(Fig 3I) from complex I, II and IV of the respiratory chain, respectively. The first two proteins are expressed from nuclear DNA, while Mtco1 is one of the 13 respiratory complex subunits encoded in the mtDNA (Friedman & Nunnari, 2014). This diminished expression may be explained by the mtDNA reduction.

In the neonatal heart, ATP is derived from glycolysis and oxidative phosphorylation, in roughly equal proportions (Lopaschuk *et al*, 1991; Piquereau & Ventura-Clapier, 2018). In the days and weeks following birth, the ATP provided by oxidative phosphorylation increases until it accounts for over 90% of the total in adult hearts (Sanchez-Diaz *et al*, 2020; Stepanov *et al*, 1997). We performed high-resolution respirometry to examine characteristics of mitochondrial oxidative phosphorylation in permeabilized cardiac muscle fibres from the left ventricle. Ventricular muscle fibres from 8-day-old Srsf3 cKO mice exhibited a significant reduction in complex I-driven respiration (Fig 3K). Respiratory chain complex I was previously reported along with complex IV to have decreased activity in a failing dog heart (Sabbah *et al*, 2016). The lower activity of complex I may contribute substantially to the heart contraction defect in Srsf3 cKO mice as the contraction/relaxation cycle in the myocardium uses approximatively 90% of the cellular ATP (Opie L.H., 1997). Mitochondrial proton leak induced by the addition of oligomycin to inhibit ATP synthase in the mitochondria energized with complex I and II substrates (*i*.*e*., malate, pyruvate, glutamate, and succinate) was not affected in the hearts of Srsf3 cKO mice, indicating an oxidative phosphorylation coupling efficiency similar as measured in Srsf3^Fl/Fl^ control hearts (Fig 3L). To determine whether the impaired fibre respiration was due to a reduction in mitochondrial content, we quantified citrate synthase activity, a biomarker that exhibits a strong association with mitochondrial content (Larsen *et al*, 2012). Citrate synthase activity was similar between conditions, suggesting an equal number of mitochondria (Fig 3M).

The folds in the inner membrane of the mitochondria are called cristae and are the major site of the complexes of the oxidative phosphorylation system. The quantity of cristae is a good index to evaluate the quality of the mitochondria (Sanchez-Diaz *et al*., 2020). In failing human and dog hearts, loss of the electron-dense matrix and marked disorganized cristae are mitochondrial abnormalities observed via transmission electron microscopy (Ferrans *et al*, 1972; Sabbah *et al*, 1992). Other abnormalities of the mitochondria can occur, such as hyperplasia, reduced organelle size, fragmentation, and disruption of the inner and outer membranes (Bonora *et al*, 2019). Transmission electron micrographs of Srsf3^Fl/Fl-βMHC-Cre^ hearts at 8 days of age revealed major cristae disorganization as well as mitochondrial matrix swelling compared to Srsf3^Fl/Fl^ hearts (Fig 3N). Change in the unit length of sarcomeres was also notable as the distance between the Z-lines was shortened in the hearts of Srsf3 cKO mice. Cardiomyocyte-specific heterogeneous nuclear ribonucleoprotein U (hnRNP U) cKO mouse model (Ye *et al*., 2015) shares similarities with the Srsf3^Fl/Fl-βMHC-Cre^ mice, namely, the rapid regression of fractional shortening and mouse death within two weeks. However, the number and size of the mitochondria as well as their alignments to the sarcomere were reduced in hnRNP U cKO mice. Those characteristics differ from the mitochondria abnormalities observed by electron microscopy and OXPHOS-related transcriptomic changes specifically identified in the heart of Srsf3 cKO mice.

These results shed new light on the unexpected relationship between Srsf3 and mitochondria in cardiomyocytes and possibly in other cell types. The production of ATP is extremely important for heart contraction, and the Srsf3 deficiency seems to impair mitochondrial ATP production by modulating the expression and splicing of mRNA. Evidently, these modifications, possibly combined with other mechanisms, such as mTOR signaling defect seen in the adult Srsf3 cKO model (Ortiz-Sanchez *et al*., 2019), provoke systolic dysfunction and lead to rapid death. Perhaps most surprising is the mitochondrial defect itself, given that the study of Srsf3 in adult mice did not display significant mitochondrial enrichment with RNA-seq coupled with gene ontology analysis. Nonetheless, it would not be the first time that something similar has been observed with a regulatory protein. The deletion of PGC-1α and β transcriptional coactivators in a mouse muscle model revealed a mitochondrial structural disorganization as well as altered expression of many genes associated with mitochondria in 2-week-old offspring mouse hearts, whereas the adult mouse model showed no major illness-related effects from the deletion (Martin *et al*, 2014). Certainly, further investigation of the impact of Srsf3 on mitochondrial dynamics in cardiac pathophysiology as well as for other cell types’ metabolism would be a promising and stimulating endeavor to pursue.

## Materials and Methods

### Animal studies

All animal procedures (Protocol # 2019-2294) complied with the NIH guidelines and were approved by the Université de Sherbrooke Institutional Committee for the Use and Care of Laboratory Animals. Srsf3-floxed mice were generously shared by Nicolas J. Webster (UCSD, USA) and bred with two different cardiomyocyte-specific Cre-overexpressing transgenic mice: βMHC-Cre (constitutive Cre recombinase expressed under *Myh7* promoter regulation in cardiomyocytes during *in utero* development) (Parsons *et al*., 2004) and αMHC-MerCreMer (tamoxifen-inducible chimeric Cre recombinase expressed under *Myh6* promoter regulation in cardiomyocytes at adulthood)(Sohal *et al*, 2001). All experiments performed with Srsf3^Fl/Fl-βMHC-Cre^ mice were performed with 5- to 15-day-old animals depending on the experiment. The primers for Srsf3, Cre and sex genotyping (Tunster, 2017) are listed in Appendix Table S3. A single intraperitoneal injection of tamoxifen (50 mg/kg, suspended in a 5% ethanol/95% peanut oil mix) was performed with 2-month-old Srsf3^Fl/Fl-αMHC-MerCreMer^ mice to efficiently invalidate the *Srsf3* gene from their cardiomyocytes. Following cardiac echography, the hearts were collected seven days post-injection to perform subsequent analysis.

### Echography analysis

Mice were anaesthetized using isoflurane (2% with 1 l/min O2, Fresenius Kabi Canada, Canada). Echocardiography was performed using a Vevo 3100 imaging system equipped with an MX400 linear array transducer (FUJIFILM VisualSonics, Canada). M-mode tracings were acquired in parasternal short-axis view of the left ventricle using the papillary muscle as a reference. Cardiac function was measured using Vevo LAB 3.1.1 software (FUJIFILM VisualSonics, Canada).

### Histology

Heart tissues were fixed via 4% paraformaldehyde (PFA), embedded in paraffin and cut into slices of 6 μm thickness. Lames were deparaffinized with 100% xylene and successive ethanol dilutions. Masson’s trichrome staining was performed on heart slices, and pictures were taken with a Hamamatsu Nanozoomer 2.0-RS with a 20x resolution. For immunofluorescence, lames were subjected to permeabilization and blockage steps with 0.5% Triton X-100 and 5% serum in PBS. Lames were stained with WGA (50 μg/ml) and IB4 (5 μg/ml) for one hour at room temperature. For immunohistology, an antigen retrieval step was performed with 10 mM sodium citrate and 0.1% Tween-20 buffer at pH 6 and incubated for 20 min at 90°C. Lames were permeabilized and blocked with 0.1% Triton X-100 and 10% serum in PBS for 30 min at room temperature. Then, the primary antibodies against Srsf3 (rabbit; Abcam, 198291) and α-actinin (mouse; Sigma, A7811 - clone EA-53) were incubated at 4°C overnight. Three washes with PBS were performed before adding DAPI and the secondary antibodies anti-rabbit Alexa 647 (Cell Signaling, 4414) and anti-mouse Alexa 488 (Cell Signaling, 4408) for 2 hours at room temperature. Images were acquired with a Leica DM4000 upright fluorescence microscope (10x and 20x objectives). Analysis of the images for cell size and capillary density measurements were performed with ImageJ software.

### Western blotting

Protein extracts were isolated from the hearts of 15-day-olds extracted with RIPA lysis buffer (50 mM Tris-HCl at pH 7.4, 150 mM NaCl, 1% Triton X-100, 1% sodium deoxycholate, 0.1% SDS, 1 mM dithiothreitol (DTT), 5 mM EDTA, and Halt Protease and Phosphatase Inhibitor Cocktail from Thermo Fisher Scientific), and the protein concentration was measured using DC Protein Assay kits (Bio-Rad). Equal quantities of proteins were resolved by SDS-PAGE gels and blotted onto polyvinylidene difluoride membranes (Millipore). Then, the membranes were blocked with 2% bovine serum albumin (BSA) before incubation with the primary and secondary antibodies. The immunoblots were visualized by using Luminata - Immobilon Crescendo Western HRP Substrate (Millipore) on a ChemiDoc MP station (Bio-Rad). Image analysis was performed using Image Lab software version 5.2 (Bio-Rad). The antibodies used for the western blots were as follows: anti-Srsf3 (Abcam, ab198291) anti-Gapdh (Cell Signaling, 2118), anti-Tim50 (Santa-Cruz, sc-393678), anti-Tom20 (Abcam, ab186735), Total OXPHOS antibodies (Abcam, ab110413), and anti-Uqcrc1 (ThermoScientific, PA5-21394).

### RT-qPCR and ASPCR

Total RNA was extracted from hearts using RNeasy Mini kits (Qiagen) according to the manufacturer’s protocol. A total of 1000 ng of total RNA was used to prepare cDNA in a 20 µL reaction with Script Reverse Transcription Supermix (Bio-Rad). qPCR was performed on technical duplicates with cDNA diluted 30x and SsoAdvanced Universal SYBR Green Supermix (Bio-Rad) via a Mastercycler ep RealPlex (Eppendorf). qPCR analysis was performed by the 2(−ΔΔCt) method, normalizing to RPL30. Splicing PCR was performed with Q5 High-Fidelity DNA Polymerase (New England Biolabs), and 5 μL of the cDNA was diluted 30x. Thirty cycles of PCR were used to produce the optimal band detectable in a 2% agarose gel. A ChemiDoc MP station (Bio-Rad) was used to reveal the ethidium bromide-stained agarose gel, and image analysis was performed using Image Lab software version 5.2 (Bio-Rad). Primers are listed in Appendix Table S3.

### Analysis of Srsf3-floxed mouse RNA-seq data

Total RNA extractions were performed on tissues using TRIzol (Invitrogen) with chloroform following the manufacturer’s protocol. Briefly, fresh mouse heart tissues were collected in 1 ml of TRIZOL and lysed using Qiagen’s TissueLyser. After the addition of chloroform, the aqueous layer was recovered, mixed with one volume of 70% ethanol and applied directly to an RNeasy Mini Kit column (Qiagen). DNAse treatment on the column and total RNA recovery were performed as per the manufacturer’s protocol. RNA integrity was assessed with an Agilent 2100 Bioanalyzer (Agilent Technologies). The mRNA was isolated from 2.5 μg of total RNA, according to the manufacturer’s recommendations (Magnetic mRNA Isolation Kit protocol from NEB, S1550S), and eluted in 17 μL. The quantity and quality of the isolated mRNAs were checked on an Agilent Pico chip. The library was built with a QIAseq™ Stranded Total RNA Lib Kit (Qiagen, 180753) from 5 μL (2 ng) of mRNA. Library quality was assessed on an Agilent DNA HS Chip and quantified by Qbit HS DNA. Libraries were then pooled and sequenced on an Illumina Nextseq 500 PE75, at the plateforme RNomique de l’Université de Sherbrooke.

### Quality control and general processing

The raw read files obtained consisted of an average of 51.1 M paired-end reads per sample with a minimum of 44.5 M reads per sample. Fastq files containing the reads were trimmed with Trimmomatic v0.38 with the parameters LEADING:30 TRAILING:30 SLIDINGWINDOW:4:15 MINLEN:50 to remove Illumina adaptors and bases with low quality (Bolger *et al*, 2014). The quality of RNA-seq reads was examined using Fastqc v0.10.1 (www.bioinformatics.babraham.ac.uk/projects/fastqc) before and after trimming.

### Validation of SRSF3 cKO efficiency

RNA-seq reads were first aligned to the Ensembl mouse genome mm.GRCm38 (release 97)(Zerbino *et al*, 2018) using STAR v2.7.1a (Dobin *et al*, 2013) with the following parameters: -- outFilterScoreMinOverLread 0.3 --outFilterMatchNminOverLread 0.3 --outFilterMultimapNmax 100 -- winAnchorMultimapNmax 100 --alignEndsProtrude 5 ConcordantPair. Differential exon usage analysis was then performed using DEXseq 1.30.0 (Anders *et al*, 2012; Reyes *et al*, 2013) with default parameters.

### Differential gene expression

Transcript abundance was first estimated using Kallisto v0.46.0 (Bray *et al*, 2016) with 100 bootstraps and the Ensembl mouse genome mm.GRCm38 (Zerbino *et al*., 2018) and then aggregated into gene abundance estimation using biomaRt v2.40.3 (Durinck *et al*, 2005; Durinck *et al*, 2009) with Ensembl annotations and summarized using tximport v1.12.3 (Soneson *et al*, 2015). Differential expression analysis was performed with DESeq2 v1.24.0 (Love *et al*, 2014). Genes with an adjusted p-value < 0.05 were considered to be differentially expressed for downstream analysis.

### Differential splicing

RNA-seq reads were first aligned using Vast-tools v2.2.2 (Tapial *et al*, 2017) to the mouse genome mm9 with default parameters. Samples of the same condition were then merged to increase read coverage following the developers’ recommendation. Differential splicing analysis was performed using Vast-tools subcommand diff (Han *et al*, 2017) with default parameters. Sashimi plots for selected events were generated using ggsashimi (Garrido-Martin *et al*, 2018).

### DNA extraction and qPCR on mitochondrial and nuclear DNA

DNA was extracted with the DNeasy Blood & Tissue Kit (Qiagen) from heart tissues and treated with RNAse A (Bio Basic, RB0471). qPCR was performed with 1 ng of DNA with the primers listed in Appendix Table S3. The relative quantity of mitochondrial DNA was obtained as a ratio between mitochondrial and nuclear DNA Ct values as previously reported (Chen *et al*, 2010a; Song *et al*, 2017).

### Permeabilized muscle fibre bundle preparation

A section of the left ventricular free wall was dissected and immediately placed in ice-cold preservation medium (BIOPS; containing 2.77 mM CaK_2_EGTA, 7.23 mM K_2_EGTA, 5.77 mM Na_2_ATP, 6.56 mM MgCl_2_·6H_2_O, 20 mM taurine, 15 mM phosphocreatine, 20 mM imidazole, 0.5 mM dithiothreitol, and 50 mM 4-morpholine-ethanesulfonic acid, pH 7.1) for mechanical separation with fine forceps under a dissecting microscope. Separated fibre bundles were incubated in BIOPS containing 5 mg/mL saponin for 30 min at 4°C and then transferred to respiration medium (MiR05; containing 0.5 mM EGTA, 3 mM MgCl_2_·6H_2_O, 20 mM taurine, 10 mM K_2_HPO_4_, 20 mM HEPES, 110 mM sucrose 60 K-lactobionate, and 1 g/L BSA, essentially fatty acid free in sterile water, pH 7.1) for three 15 min washes at 4° C (Kanaan *et al*, 2018).

### Mitochondrial respiratory capacity (JO_**2**_)

Respiration of permeabilized muscle fibre bundles was measured by high-resolution respirometry (Oxygraph-2k modular system, OROBOROS Instruments, Austria) in MiR05 supplemented with creatine (20 mM) and blebbistatin (5 μM) at 37°C, and oxygen concentrations were maintained between 450 and 250 μM in duplicate and analysed using Datlab software (OROBOROS Instruments Corp., Innsbruck, Austria). The respirometry protocol consisted of the sequential addition of 2 mM malate (M) and 5 mM pyruvate (P) (adenylate free leak respiration; Complex I leak or CI_leak_), followed by the addition of 4 mM ADP (D) and 10 mM glutamate (G) (complex I-supported respiration; CI_ADP_). Then, maximal oxidative phosphorylation (OXPHOS) capacity with convergent electron flow from complex I and complex II was reached with 10 mM succinate (S) (CI+CII_ADP_). ATP synthesis was then inhibited using 2.5 μM oligomycin to inhibit ATP synthase and induce non-phosphorylating leak (CI+CII_Leak_). Finally, non-mitochondrial respiration was measured after complex III inhibition by 5 μM antimycin. All reported respiration measures are corrected for non-mitochondrial respiration. Outer mitochondrial membrane integrity was confirmed with the addition of 10 μM cytochrome c.

### Citrate synthase activity

Frozen left ventricular tissue was homogenized using a bead mill ruptor (Fisherbrand Bead Mill 24 Homogenizer) in ice-cold RIPA buffer (20 mM Tris-HCl, pH 7.5, 150 mM NaCl, 1 mM Na_2_EDTA, 1 mM EGTA, 1% NP-40, 1% Na deoxycholate, 2.5 mM Na pyrophosphate, 1 mM β-glycerolphosphate, 1 mM Na_3_VO_4_) supplemented with protease and phosphatase inhibitors (Sigma Aldrich, P8340 and P5726). Cellular debris were removed by centrifugation at 4°C for 10 min at 14,000g. Protein concentrations were determined using a BCA kit according to the manufacturer’s protocol (Pierce BCA Protein Assay Kit, Thermo Fisher Scientific, Rockford, IL, USA). Citrate synthase (CS) activity was determined by measuring the change in absorbance at 412 nm in 50 mM Tris-HCl buffer (pH 8.0) containing 0.2 mM 5,5-dithiobis(2-nitrobenzoic acid) (DNTB), 0.1 mM acetyl-coA, and 0.25 mM oxaloacetate using the BioTek Synergy Mx Microplate Reader at 25°C. Specific activity of the CS was calculated using the extinction coefficient of DNTB of 13.6 mM^-1^cm^-1^ and normalized to protein content.

### Transmission electron microscopy

Heart tissues were fixed in 2.5% glutaraldehyde in 0.1 M sodium cacodylate buffer (pH 7.4). The sample was then fixed in 1% OsO4 in 0.1 M cacodylate buffer (pH 7.5) and thoroughly dehydrated in ethanol solutions. After dehydration, tissues were infiltrated with epoxy-embedded medium and cut into slices. The slices were affixed on a copper grid of 300 mesh. Pictures were acquired with a transmission electron microscope (magnification: 7000X and 25 000X).

### Statistical analyses

Data are expressed as the means and standard deviations. Statistical analysis was performed with GraphPad Prism 6 software to determine the significant difference between groups by either the T-test with the Mann-Whitney post hoc test (comparison of 2 groups: Srsf3^Fl/Fl^ and Srsf3^Fl/Fl-βMHC-Cre^) or one-way ANOVA with Tukey’s post hoc test (comparison of 3 groups: Srsf3^Fl/Fl^, Srsf3^Fl/+-βMHC-cre^, and Srsf3^Fl/Fl-βMHC-Cre^), and p-values < 0.05 were considered significant. Overlap for differentially expressed genes and differentially spliced events between adult and 5-day-old mice was calculated, and statistical significance was determined by a hypergeometric test using SciPy v1.4.1.

## Data availability

All the RNA-seq data are available from GEO (GSE151168; https://www.ncbi.nlm.nih.gov/geo/query/acc.cgi?acc=GSE151168).

## Acknowledgements

We are grateful to the Université de Sherbrooke RNomics Platform (Elvy Lapointe, Philippe Thibault) for insights into RNA-seq experiments. We thank Pr Jean-Luc Parent for sharing antibodies. This work was supported by grants from the Canadian Institute of Health Research (grant number 301131 to M.A.-M., and FDN 143278 to M-E.H.) and a Centre de Recherche du CHUS operational grant to D.P.B., training awards from Fonds de Recherche du Québec – Santé (FRQS) and Canadian Institutes of Health Research to A.-A.D., a training award from the Faculté de Médecine et des Sciences de la Santé - Domenico-Regoli to L.D., a training award from CIHR Frederick Banting and Charles Best graduate scholarship to H.G., and FRQS-Junior 1 and Heart and Stroke Foundation of Canada (HSFC) New Investigator awards, both awarded to M.A.-M.

## Author contributions

M.A.-M. supervised the study and obtained and managed funding for the project. A-A.D. and M.A-M. conceptualized the experiments. A-A.D., L.D., D.Z., H.G., C.P., M.-E.H., D.P.B., M.S.S., and M.A-M. performed the experiments and/or analysed the results. A-A.D. and M.A-M. wrote the article. All authors reviewed the manuscript.

## Conflict of interest

The authors declare that they have no conflicts of interest.

## Appendix figure and table legends

**Table S1.** Echography analyses. Analysis of the parasternal long axis (PLAX) and short axis (SAX) echocardiographic views obtained from 15-day-old control and Srsf3 cKO mice. The following left ventricle (LV) parameters represent the LV end-systolic volume (LVESV), LV end-diastolic volume (LVEDV), LV end-diastolic anterior wall thickness (LVAWd), and LV end-diastolic posterior wall thickness (LVPWd). n=6 for each genotype, *p-value compared to Srsf3^Fl/Fl^ mice and #p-value compared to Srsf3^Fl/+-βMHC-Cre^ mice.

|  |  | Srsf3<sup>Fl/Fl</sup> |  | Srsf3<sup>Fl/+βMHC-Cre</sup> |  | Srsf3<sup>Fl/Fl-βMHC-Cre</sup> |  |  |
| --- | --- | --- | --- | --- | --- | --- | --- | --- |
| PLAX | Ejection Fraction (%) | 59.2 | ±6.3 | 59.6 | ±6.4 | 29.8 | ±10.6 | *<0.0001 and #<0.0001 |
|  | Heart Rate (beats/min) | 431.5 | ±23.4 | 401.9 | ±32.9 | 346.0 | ±57.0 | * 0.0071 and # 0.0379 |
|  | Cardiac Output (mL/min) | 8.6 | ±2.5 | 6.6 | ±2.2 | 5.0 | ±1.8 | *0.0255 |
|  | LVESV (μL) | 13.7 | ±4.6 | 11.0 | ±2.6 | 42.2 | ±28.4 | *0.0249 and # 0.0135 |
|  | LVEDV (μL) | 33.5 | ±8.7 | 27.3 | ±6.5 | 57.3 | ±31.3 | # 0.04 |
|  | Stroke Volume (μL) | 19.8 | ±5.2 | 16.3 | ±5.0 | 15.1 | ±5.7 |  |
| SAX | Fractional Shortening (%) | 28.7 | ±3.3 | 27.5 | ±5.1 | 16.4 | ±6.1 | * 0.0017 and # 0.0039 |
|  | LV Mass Corrected (mg) | 47.2 | ±13.8 | 44.6 | ±12.9 | 62.4 | ±34.4 |  |
|  | LVAWd (mm) | 0.6 | ±0.1 | 0.7 | ±0.2 | 0.5 | ±0.1 |  |
|  | LVPWd (mm) | 0.6 | ±0.1 | 0.5 | ±0.2 | 0.6 | ±0.2 |  |

**Figure S1.**
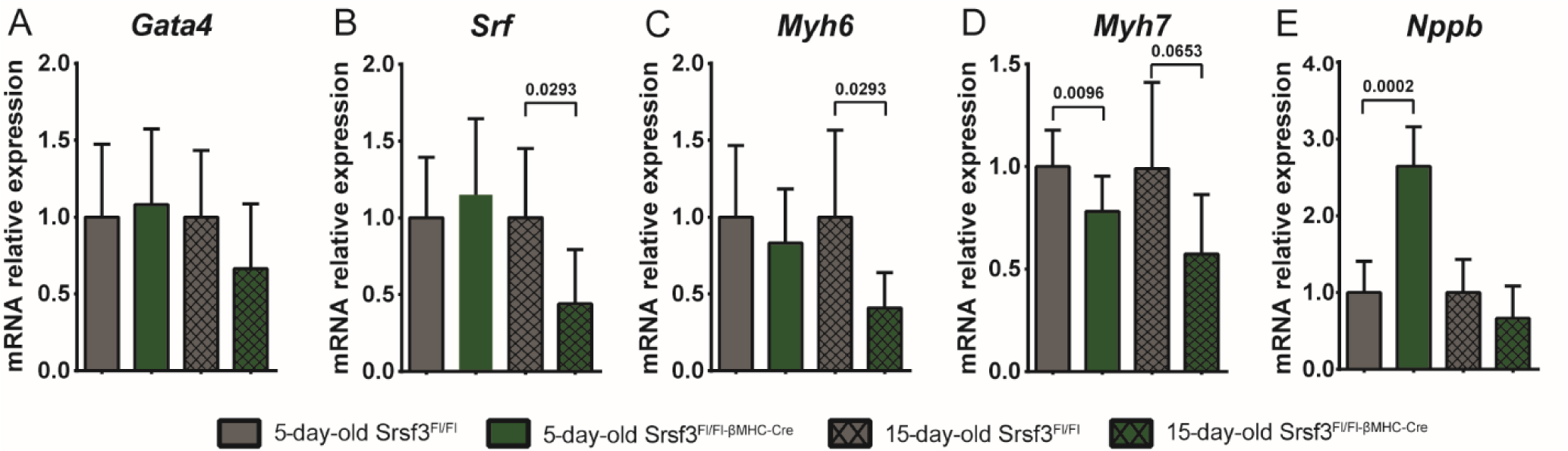
mRNA diminution of Srsf3 and hypertrophic gene panel. Analysis of the mRNA by RT-qPCR of heart samples in 5- and 15-day-old control Srsf3^Fl/Fl^ and Srsf3^Fl/Fl-βMHC-Cre^ mice for **A)** *Gata4*, **B)** *Srf*, **C)** *Myh6*, **D)** *Myh7*, and **E)** *Nppb*.

**Figure S2.**
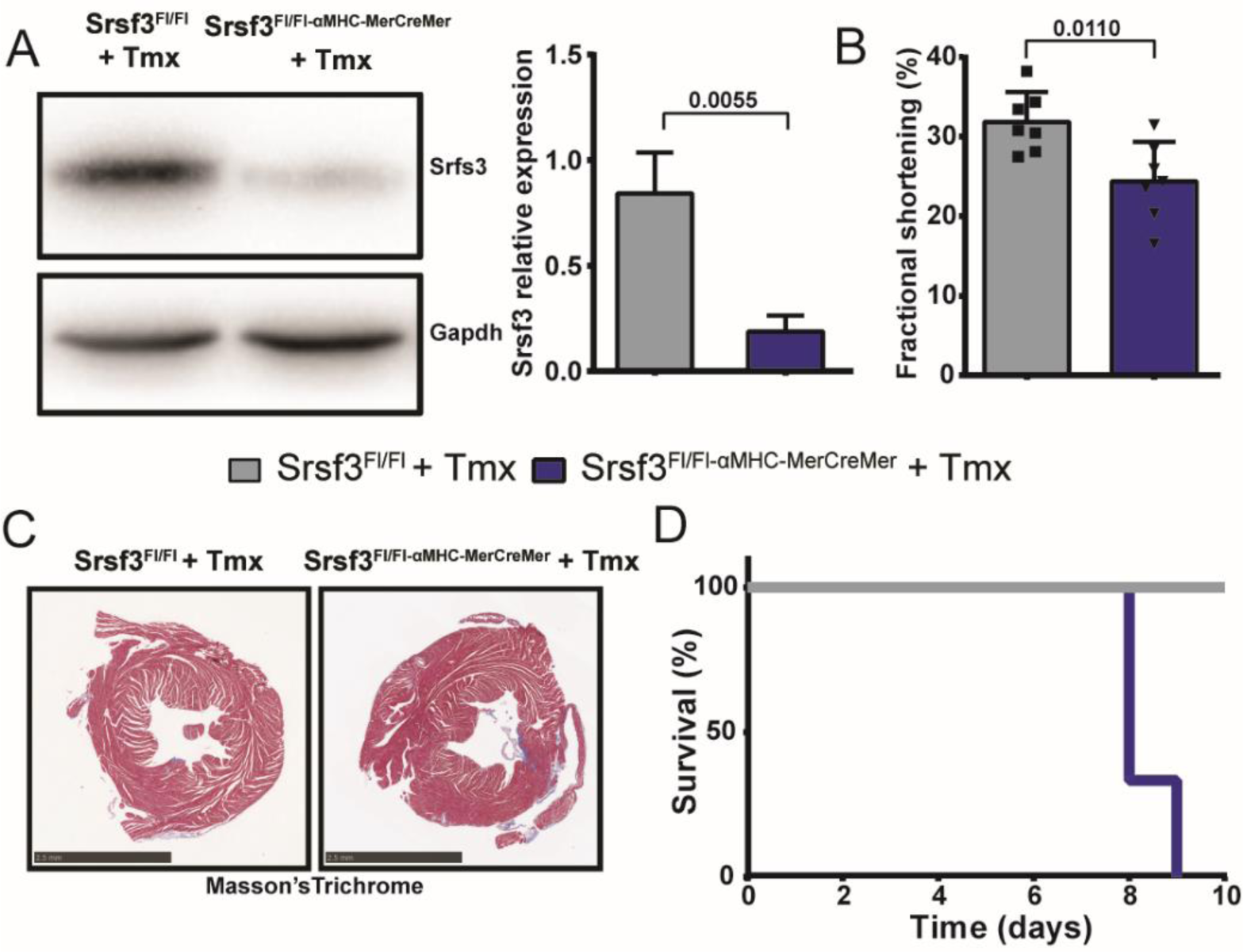
Deletion of Srsf3 in cardiomyocytes in adult mice diminishes the cardiac function and survival. **A)** Western blot with the graphic quantification of SRSF3 from heart samples in two mouse genotypes (Srsf3^Fl/Fl^ and Srsf3^Fl/Fl-αMHC-Mer-Cre-Mer^) seven days after tamoxifen injection. **B)** Heart contractility analysed by fractional shortening in the presence or absence of Srsf3 seven days after tamoxifen injection. **C)** Heart transverse slices stained by Masson’s trichrome (scale bar: 2.5 mm). **D)** Survival curve after the injection of tamoxifen in the Srsf3^Fl/Fl^ and Srsf3^Fl/Fl-αMHC-Mer-Cre-Mer^ mice (n = 3 for each group).

**Table S2.**
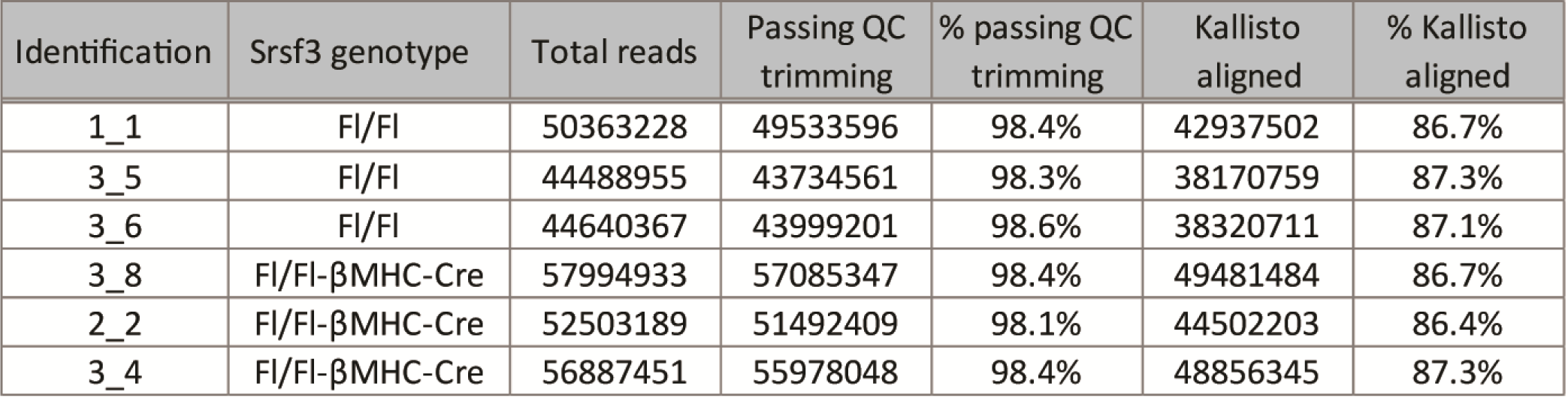
List of reads from RNA-seq analysis. Table showing the read numbers and percentage of reads kept from the Illumina sequencing and after the trimming and the alignment analysis for all the samples (Srsf3^Fl/Fl^ or Srsf3^Fl/Fl-βMHC-Cre^; n=3 for either group).

**Table S3.**
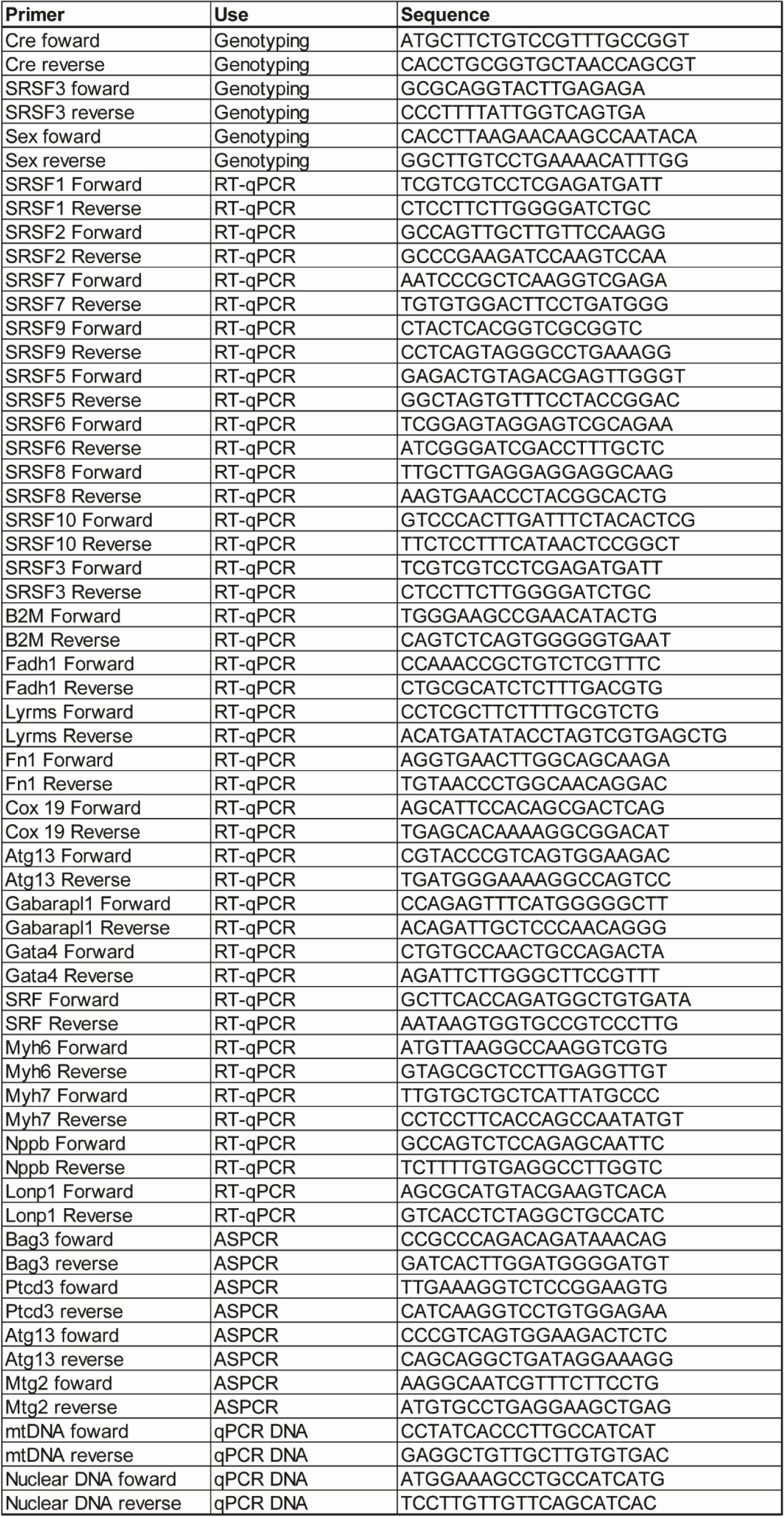
List of primers. Primers used for the RT-qPCR, ASPCR, mouse genotyping, and DNA qPCR experiments.

**Figure S3.**
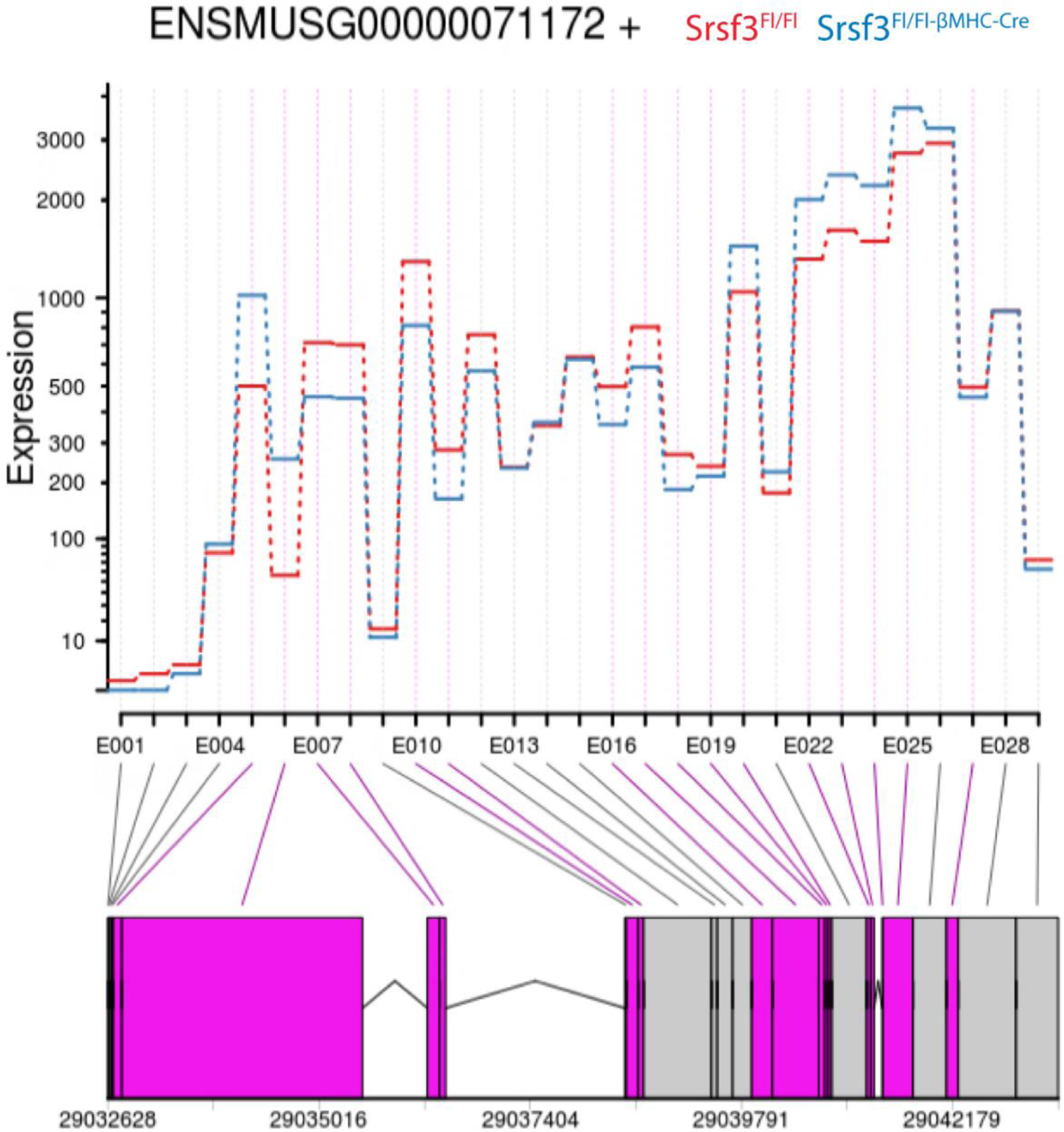
Quantification of the Srsf3 knockout from RNA-seq heart samples. Analysis of the RNA expression of Srsf3 in the control Srsf3^Fl/Fl^ (red) and Srsf3^Fl/Fl-βMHC-Cre^ (blue) mice. The flox sequences are positioned around exons 2 and 3, which correspond to sections E007 to E010 by DEXseq. The purple boxes below represent significant expression changes between the conditions. Overall, the cardiac expression level of the Srsf3 transcript from Srsf3^Fl/Fl-βMHC-Cre^ mice was equivalent to 57% of that detected from the control Srsf3^Fl/Fl^ mice.

**Figure S4.**
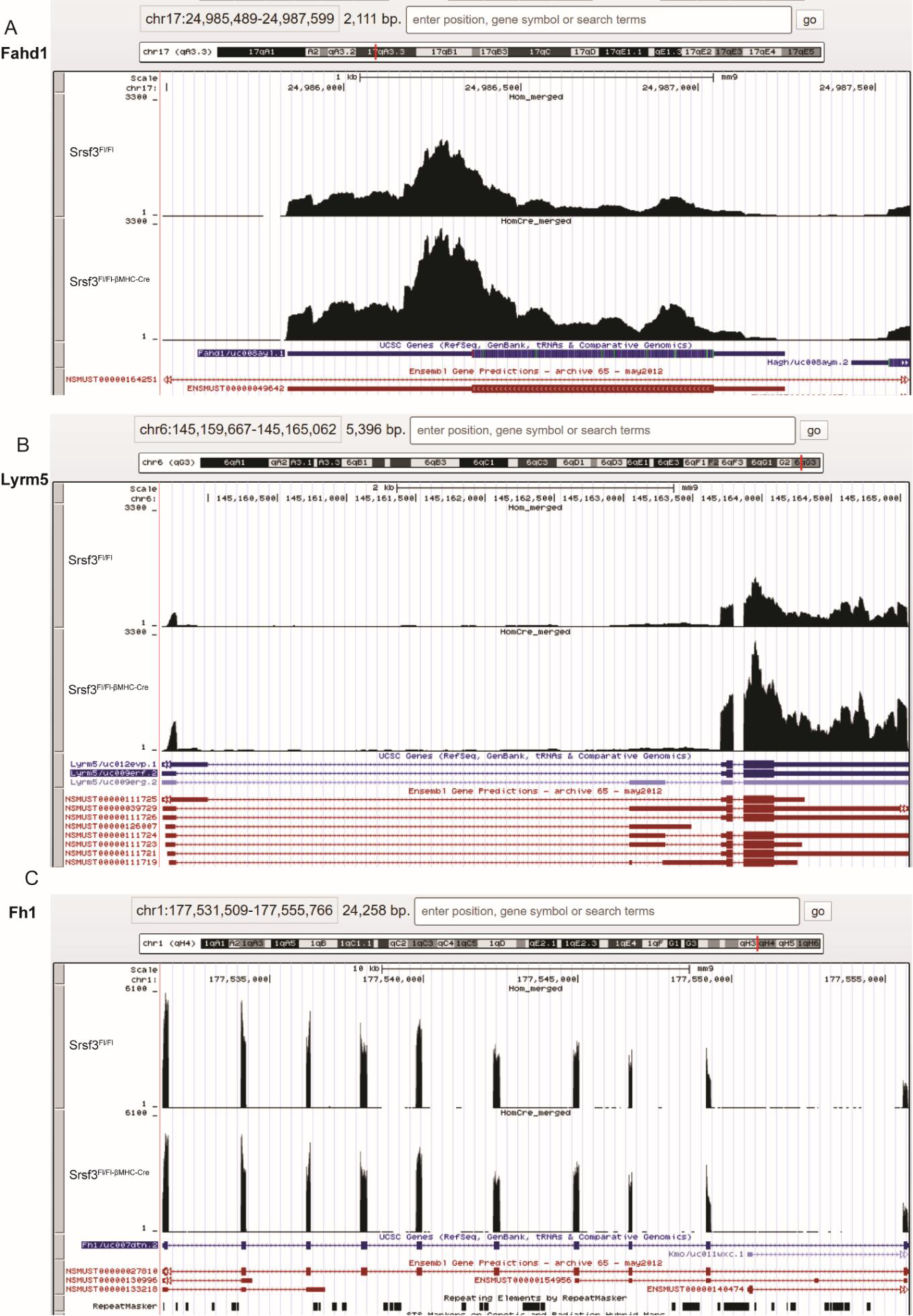
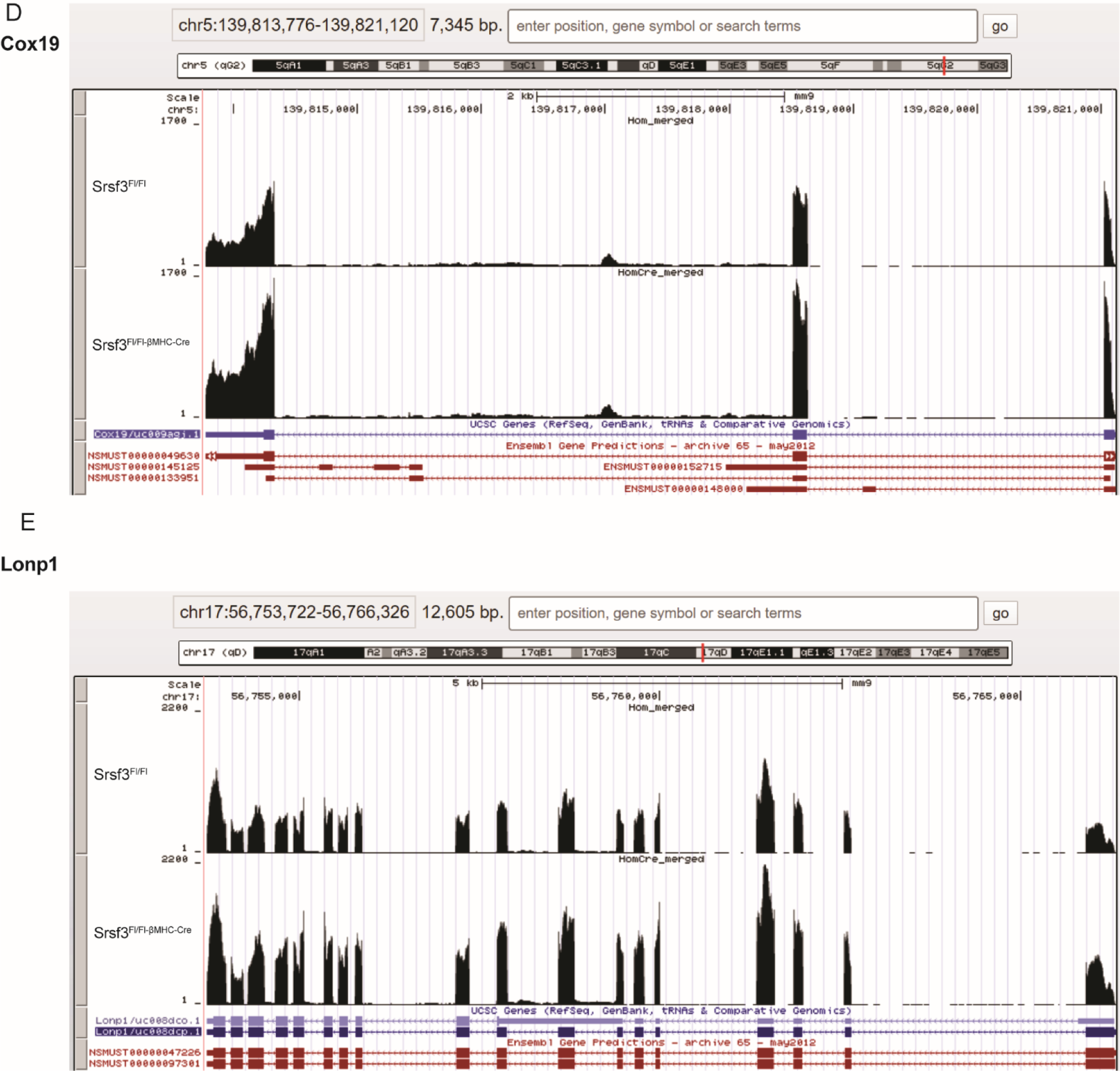
The absence of Srsf3 in the heart impacts the expression of mRNA with a mitochondrial function. Screenshots of the UCSC genome browser displaying the number of reads aligned to each gene of the mRNA analysed for their biological function linked to the mitochondria, *i*.*e*., **A)** Fadh1, **B)** Lyrm5, **C)** Fh1, **D)** Cox19, **E)** Lonp1, **F)** Atg13, and **G)** Gabarapl1.

**Figure S5.**
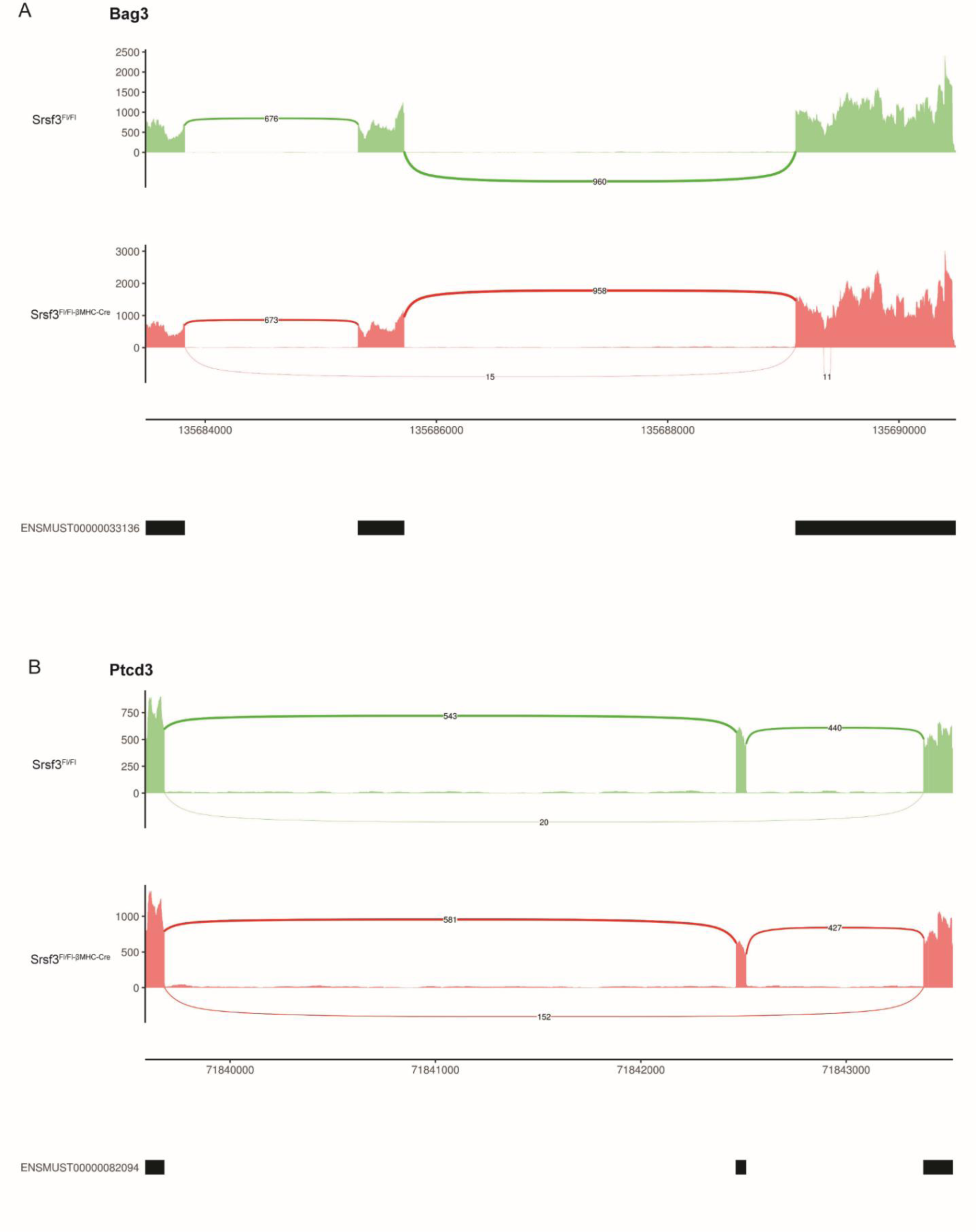
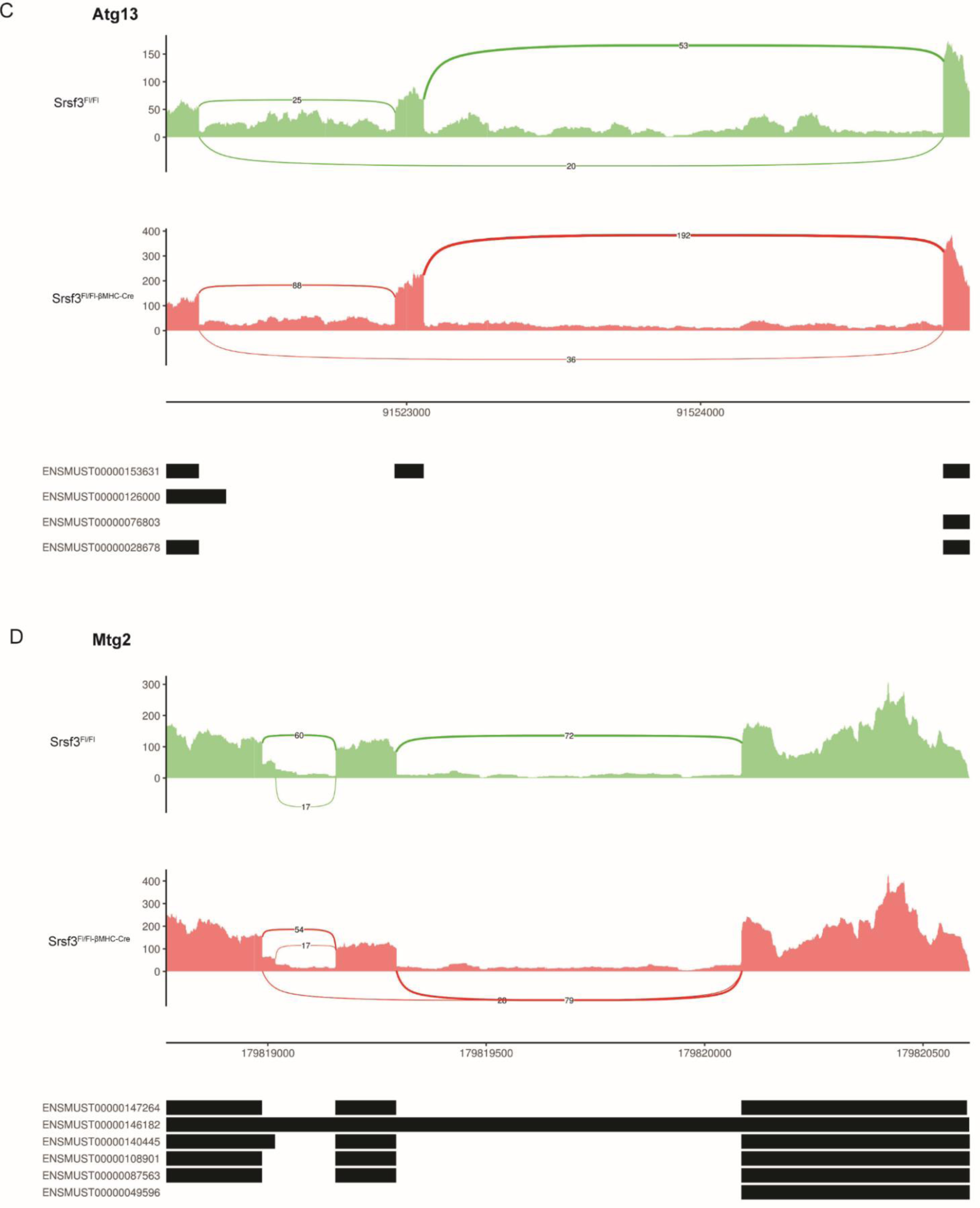
Alternative splicing of some mRNAs with mitochondrial function in the absence of Srsf3 in cardiomyocytes. Sashimi plots representing an alternative splicing event of different mRNAs with a biological function related to the mitochondria, *i*.*e*., **A)** Bag3, **B)** Ptcd3, **C)** Atg13, and **D)** Mtg2. Numbers of aligned genomic reads are converted into read density plotted as the y-axis value, while junction reads are plotted as arcs whose width is proportional to the number of reads aligned to the junction spanning the exons connected by the arc. Exons from transcripts annotated by Ensembl are shown as black boxes below the graph for comparison.

## Notes

https://www.ncbi.nlm.nih.gov/geo/query/acc.cgi?acc=GSE151168

### Competing Interest Statement

The authors have declared no competing interest.

